# A comprehensive investigation of colorectal cancer progression, from the early to late-stage, a systems biology approach

**DOI:** 10.1101/2020.10.24.353292

**Authors:** Mohammad Ghorbani, Yazdan Asgari

## Abstract

Colorectal cancer is a widespread malignancy with a concerning mortality rate. It could be curable at the first stages, but the progress of the disease and reaching to the stage-4 could make shift the treatments from curative to palliative. In this stage, the survival rate is meager, and therapy options are limited. The question is, what are the hallmarks of this stage and what genes are involved? What mechanism and pathways could drive such a malign shift from stage-1 to stage-4? In this study, first we identified the core modules for both the stage-1 and stage-4 which four of them have a significant role in stage-1 and two of them have a role in stage-4. Then we investigated the gene ontology and hallmarks analysis for each stage. According to the results, the immune-related process, especially interferon-gamma, impacts stage-1 in colorectal cancer. Concerning stage-4, extracellular matrix ontologies, and metastatic hallmarks are in charge. At last, we performed a differentially expressed gene analysis of stage-4 vs. stage-1 and analyzed their pathways which reasonably undergone a hypo/hyperactivity or being abnormally regulated through the cancer progression. We found that lncRNA in canonical WNT signaling and colon cancer has the most significant pathways, followed by WNT signaling, which means that these pathways may be the driver for the development from early-stage to late-stage. Of these lncRNAs, we had two upregulated kind, H19, and HOTAIR, which both can be involved and mediate metastasis and invasion in colorectal cancer.

## Introduction

From submocusa it arose, growing to beyond the inner layer; no invasion, manageable and curable; this is what you can call an early-stage(stage-1) of colorectal cancer [1]. Usually, it needs imaging and colonoscopy to diagnose [2]. The primary strategy at this point is resection of the tumor [3]. Even if therapy has become successful, constant surveillance must continue to secure that the tumor never comes back, yet there is still a risk of recurrence [4]. One of the reasons for this ominous turn of events and progress of the disease is Micrometastasis in stage-1/2, which enables tumor cells to reach the nearby lymph node and hide from the available detection method. These cells could be responsible for later relapse [5]. Still, the survival rate for early-stage is about 93.2% percent. As the disease progress, survival goes down with it; Stage-2 has a survival rate of 85 to 70 percent, Stage three is about 44 percent, and finally, only eight percent of late-stage(stage-4) patients could pass the five-year survival [6]. In Stage-2, cancer might be grown into the wall of the bowl, to the adjacent tissues [7]. Surgery and adjuvant chemotherapy and different kinds of combination therapy have been considered as an option at this point. Despite all of this, the risk of recurrence in this stage is highly increased [8]. By the Stage-3, tumor growth further to bowel wall and even into the nearby organs. It has already entered lymph nodes [7], and it is safe to say that it shows far aggressive behavior. At this level, recurrence is more common, the survival rate drops drastically, and managing tumor becomes harder until the late-stage occurs [9]. Stage-4 of colorectal cancer begins with the spread throughout the body [10]. Usually liver is the first place of the invasion which followed by the lung, bones, and brain; this is what we can call advanced colorectal cancer with a grave prognosis [11]. Late-stage usually response partially to treatment, and of course, at this stage, relapse and recurrence are a prevalent phenomenon [12, 13]. Indeed, one of the hallmarks of late-stage is the invasion [14].

So, there is an urgent need to understand what could contribute to this transformation from stage-1 to stage-4. What pathway involves and what modules are in charge? Systems biology could provide an answer for these questions by analyzing high-throughput data. It enables us to infer from the expression profile, so what modules are involving in different Stages of the disease and what pathways make the difference. It expands our insight and opens new possibilities to treat and handle the diseases [15]. Since we must consider the worst scenario, we should have a good idea about the mechanism of early and late stages of colorectal cancer and have a piece of proper knowledge about what could make this progress happens. If there is a possibility to stop this transformation from stage-1 to stage-4, it must be considered and taken into effect.

In this investigation, we analyze the most valuable modules by using the weighted co-expression network analysis (WGCNA). Based on the integrative results of WGCNA modules, we identified the essential genes in each stage. Then we conducted a thorough Gene Ontology (GO), Hallmark’s analysis for both the early and the late-stage of colorectal cancer. To identify the accountable pathway for this process, we performed a differentially expressed gene(DEG) analysis between stage-4 and stage-1. The outcome of our DEG investigation went through a pathway analysis in order to identify the responsible genes and pathways.

## Materials and methods

### Research design

Our main question was the primary features of the first and late stages of colorectal cancer and how this progression occurs. We used the system biology approach to answer these questions. We had found primary modules in each stage, using hallmarks and gene ontology analysis, then we identified the main features. Nonetheless, what could bring such a contrast? We conducted a differentially expressed gene analysis to answer this question, and the outcome underwent a pathway analysis. Our conclusion was based on The main pathways and their relationship with their respected genes.(Figure 1)

**Figure 1.**
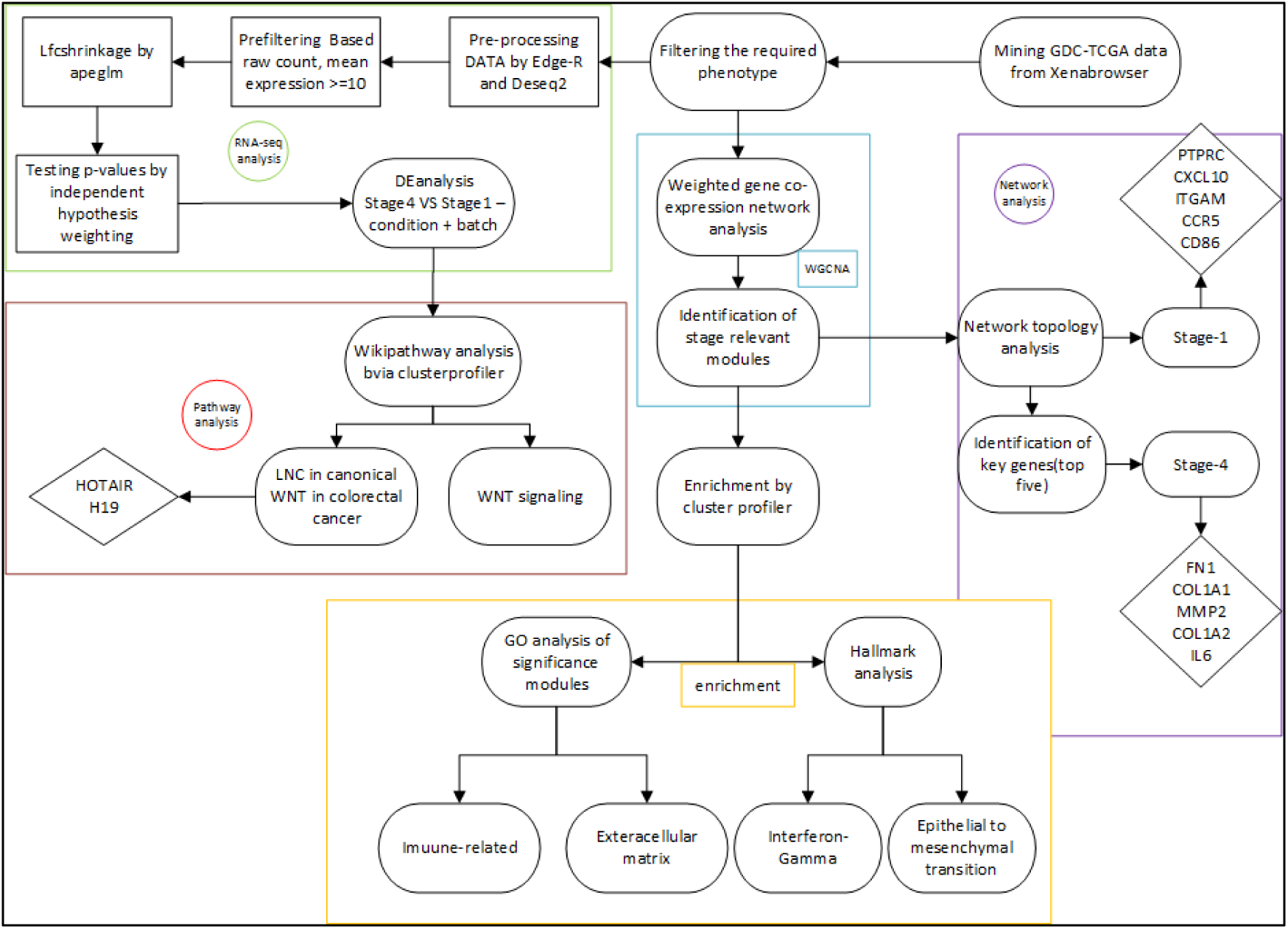
Workflow of the **s**tudy

### Adjusting phenotype to different stages

First, we obtained phenotype data from xenabrowser, a TCGA tool. Data comes from “GDC TCGA COAD” dataset 2019 version [16]. In order to filter typical phenotypes based on our purpose, which was identifying functional module and key-genes in stage-4 and stage-1 of colon cancer, we set the parameter accordingly: sample type on “primary tumor” because we need to explore stage-4 and stage-1. Hence the healthy solid tissues were unrelated. Primary diagnosis on “Adenocarcinoma”, and disease type was on “Adenomas and adenocarcinomas” because it comprises the most cases of colon cancer. Regarding tumor stages, Stage-I and stage-Ia considered as stage-1 with 68 samples, and stages-IV/IVa/IVb considered as stage-4 with 56 samples. We also included Tumor Node Metastasis (TNM) data since it could be directly applied to colorectal cancer and can define different stages in this disease. Then we matched the identified sample id from phenotype sheet to count and FPKM data for further analysis (Supp1).

### Weighted Gene Co-expression Network Analysis

Weighted Gene Co-expression Network Analysis (WGCNA) is an R package that can find clusters of highly correlated genes that are valuable in a biological context and relate them to a set of distinct traits, and this could directly guide you to hub-genes, hub-modules, and even significance pathways concerning to that specific trait [17]. First, we pre-processed FPKM data, related to early and late stages of colorectal cancer, and removing outlier rows with zero median expression, since they do not have any biological value. Then we chose the top quartile of our network by variance. This process conducted by R version 3.6.3 [18]. Mentioned pre-processed data again, going through sample clustering via WGCNA to trim outlier samples. For this analysis, we chose the Step-by-step network construction method for the identification of correlated modules. After drawing scale-free topology and choosing seven as soft power, the process of building the adjacency matrix started, which then transformed into the topological overlap matrix and then to its respective similarity (TOMsimilarity) and dissimilarityTOM matrix. Finally, a cluster dendrogram established based on TOM-dissimilarity and cutting of the eigengenes modules tree. To discover the essential genes, we merged all modules which met our given criteria (significance>0.1 and P-value <0.1). For stage-4, midnight blue and black modules were statistically meaningful, and in stage-1, there were red, green yellow, grey60, and magenta.

### Topological Analysis

Obtained integrated modules from late-stage and early-stage were imported to Cytoscape to assemble a Protein-Protein Interaction (PPI) network by STRING database (Search Tool for the Retrieval of Interacting Genes/Proteins) [19, 20]. Confidence cutoff was set on 0.4, and the maximum additional interaction was zero. Obtained PPI network undergoes a centrality-based analysis via CytoNCA, a Cytoscape plugin; CytoNCA is a centrality ranking method and investigates an assigned network by eight centralities method [21].

### Enrichment

To find a significant pathway in the most correlated modules, we ran a GO over-representation analysis (ORA) through an R package called clusterprofiler [22–24]. Then we use Msigdb analysis (another clusterprofiler features), which allows us to perform hallmarks analysis [13, 25]. Additionally, to find a significantly enriched pathway in differentially expressed genes, we ran a gene set enrichment analysis (GSEA) by clusterprofiler through wikipathway database in a sort of pre-ranked list. We used fold change as a weight for enrichment. The criteria for these analyses was log2foldchange>=1 and adjusted-value <=0.05. Besides, we used enrichplot (an R package) to draw plots and figures to be more visually illustrative [26].

### Identifying differentially expressed genes

First, we downloaded raw count data from xenabrowser, GDC TCGA COAD dataset. However, data was in the form of log2 (count+1) and needed to be transformed to the original count table (count without log) before using it as raw material for DEanalysis packages in R. According to phenotype choice, stage-1 consider as control by 68 samples. and stage-4 consider as the case by 56 Samples. Then we used EdgeR (an R package) to get annotations through “org.Hs.eg.db” only to keep rows that have Entrez Gene ID [27, 28]. In the next step, we used Deseq2 (another R package) to keep rows that meet our criteria for counts expression rowsums>=10 [29]. Moreover, LFC shrinkage applied by apeglm method and Independent hypothesis weighting estimates weighs for each P-value through the Benjamini-Hochberg method [30, 31]. We only considered genes statistically meaningful if their Adjusted P-value was less than or equal to 0.05. all computational analyses were performed by R (v 3.6.3) and Rstudio [32] in a windows10 OS.

## Results

### Identification of primary-modules

Our sample clustering detected and removed 19 outliers from our first collection of samples. Scale-free topology identified seven as an optimal power (Figure 2a). We cut the eigengenes module tree by 0.30 to correlate to 0.70 similarity. Based on our inquiry 17 modules found in the network that two of them can be correlated to stage-4 (black and midnight blue) and four of them met the criteria to be considered as significant in stage-1 category (green yellow, red, grey60, and magenta) (Figures 2b, 2c). Also, Midnight blue Module appeared to correlate to N and T stages in T.N.M staging system (Figures 2b, 2c). But the Black module seemed to be more associated with M stage also, the difference was very marginal (Figures 2b, 2c). Then we integrated all statistically meaningful modules in each stage in order to make a gene ontology analysis. In this analysis, our networks considered as a signed network since both negatively and positively correlated genes considered important in the related modules.

**Figure 2:**
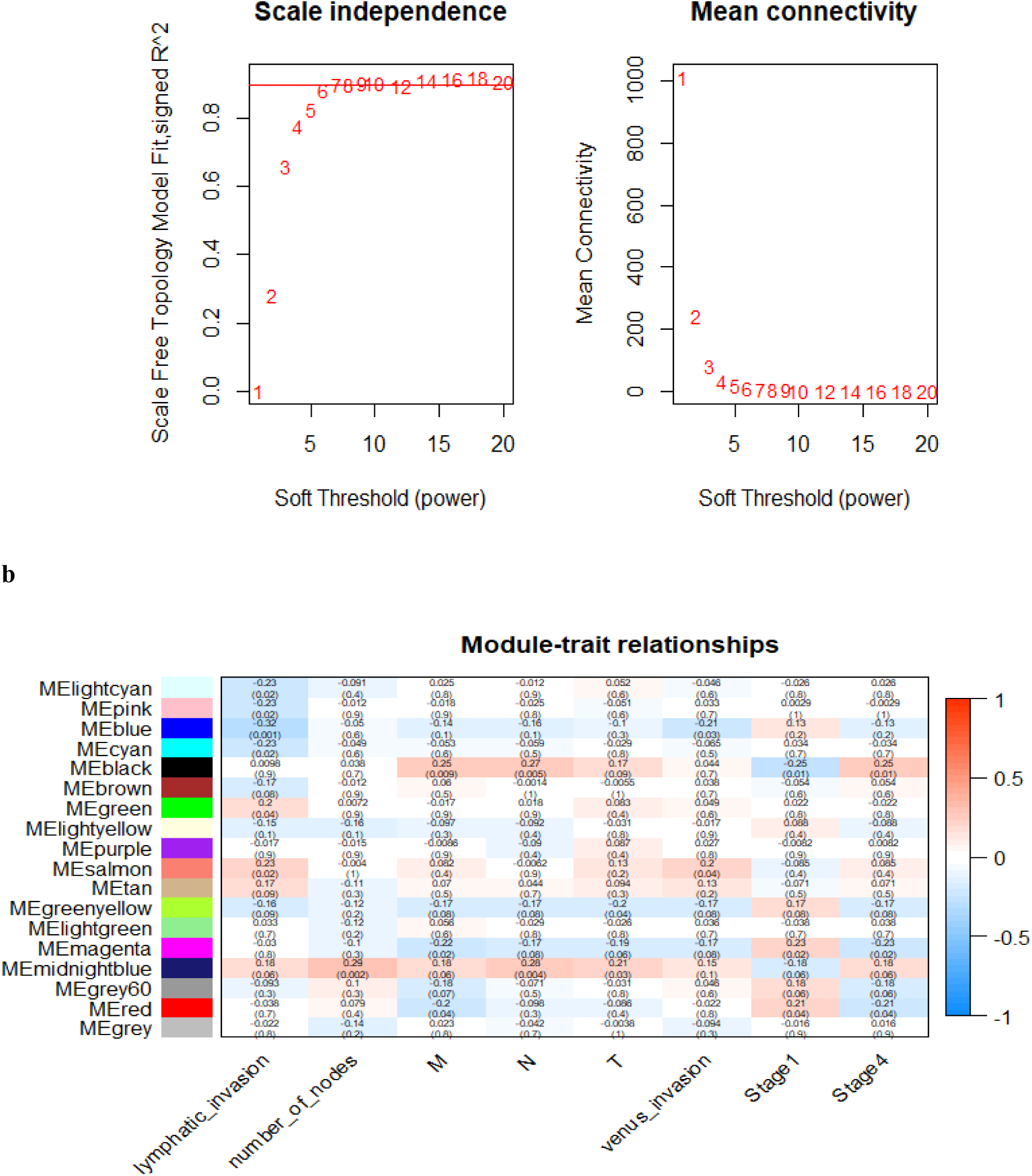
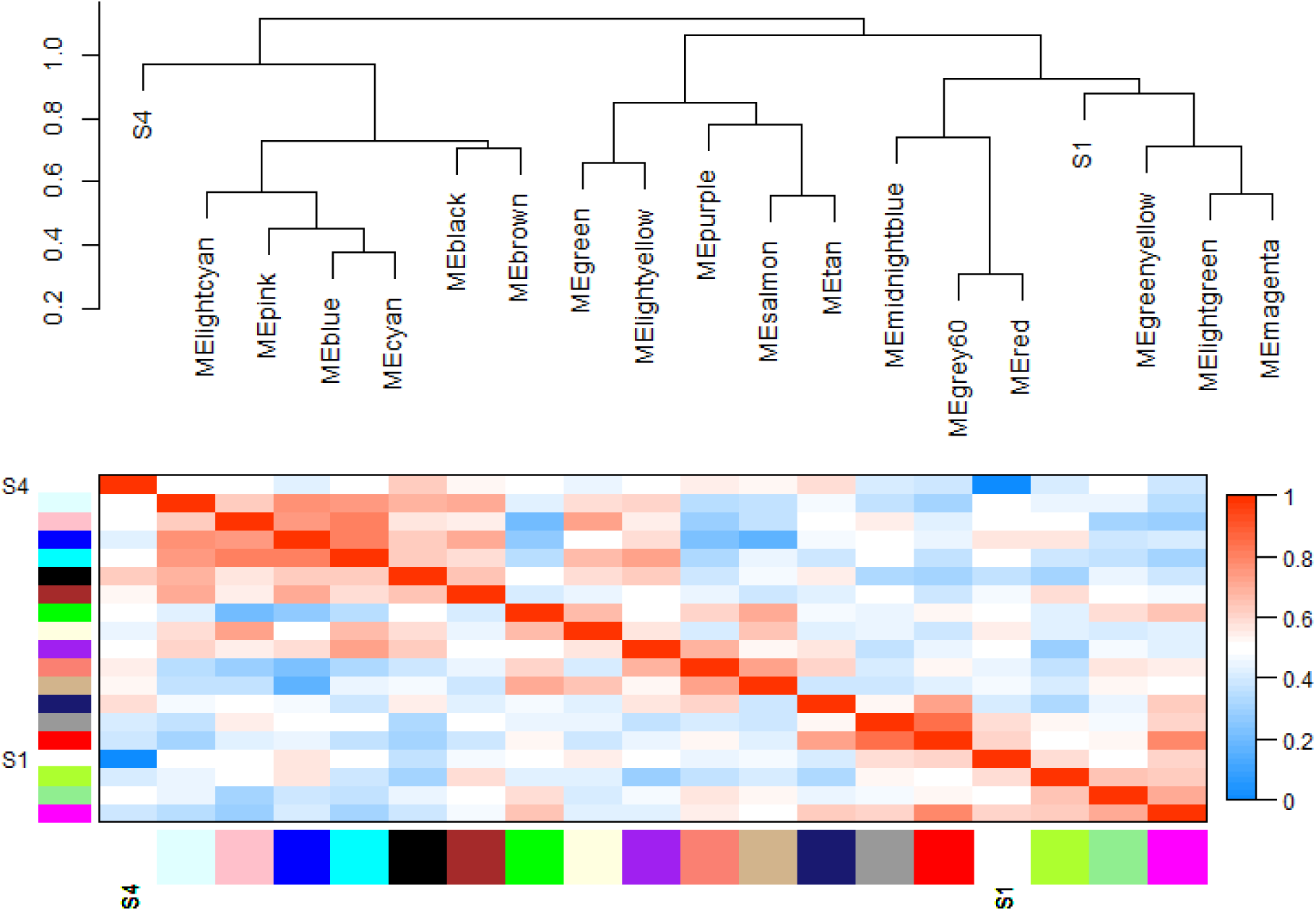
**a)** Scale-free topology to obtain the optimal soft power, **b)** Module trait relationship in which black and midnight blue modules came as significant for stage-4, and green yellow, red, grey60, and magenta came as significance for stage-1, **c)** Correlation between modules and stages using plotEigengeneNetworks.

### Module Enrichment

To get a deep insight into the functionality of the modules, we performed a GO over-representation analysis via clusterprofiler. GO analysis implemented in three categories: biological pathway, cellular component, and molecular function [22].

In stage-1, four modules met our criteria for GO enrichment (green yellow, magenta, red, and grey60). We integrated all genes from those modules and conducted a GO analysis in three categories. Biological pathway mostly pointed out immune-related process (Figure 3). Furthermore, cellular component comprised of immune system features, like immunoglobin complex and MHC classes. Molecular function was considered chemokine and cytokine activity, antigen binding, and other immune like trends as significance. Since stage-4 only had two statistically notable modules, we merged these two as one unified block (black and midnight blue). The GO results from cellular component, biological pathway, and molecular function for this specific block was very consistent. In biological pathway, extracellular matrix organization and extracellular structure organization by far had the most meaningful adjusted-p-value, q-value, and gene ratio. However, the majority of results involved in matrix, collagen, and adhesion processes (Figure 3). In cellular component category, collagen-containing extracellular matrix had the best score by far, and most of the top ten outcomes led us to the extracellular matrix and cell adhesion. There was also evident in the top ten molecular function.

We also run an MSigDb analysis in order to find hallmarks for each stage [25]. According to stage-1 hallmarks, “interferon-gamma (IFN-Y) response” had the most significance adjusted p-value and q-value (Figure 4). Results of the stage-4 modules (black and midnight blue) hallmarks were a closure to our GO analysis (Figure 4). hallmarks analysis had a solid answer to our GO results, and that was Epithelial to mesenchymal transition, which is very consistent with late-stage behavior and proved by many experimental and clinical studies [33, 34].

**Figure 3:**
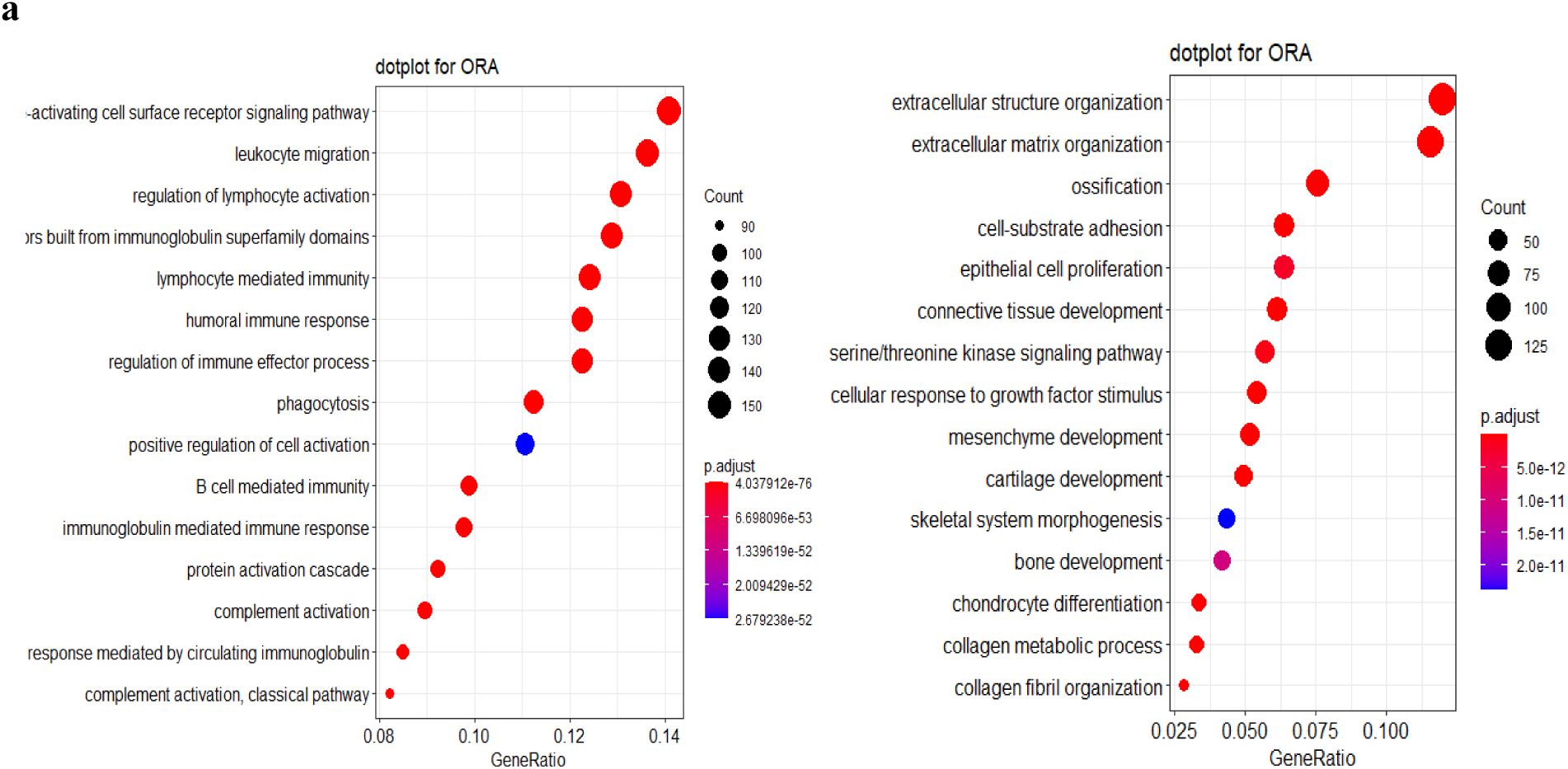
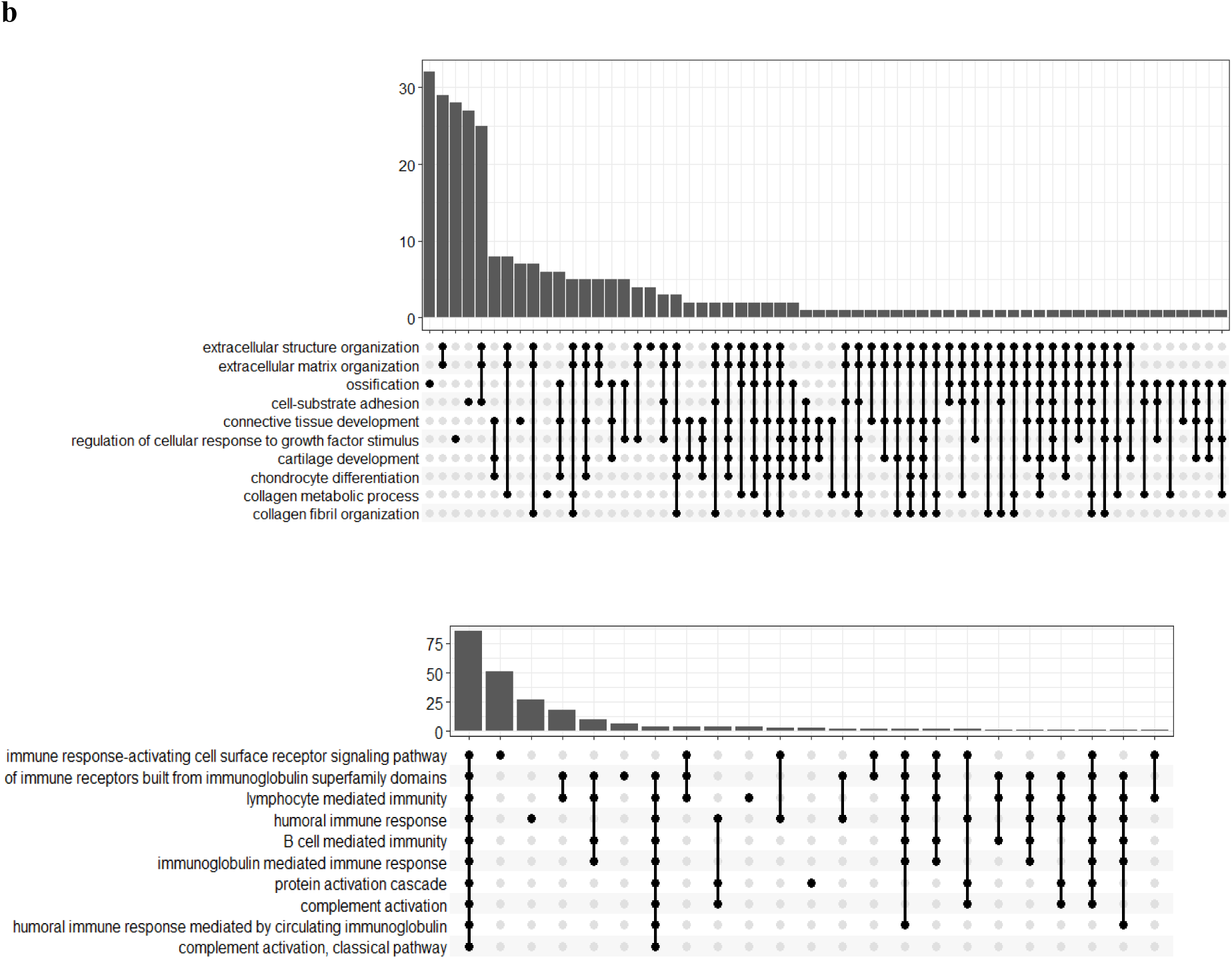
**a)** A dot plot of GO biological pathways, terms are categorized based on their gene ratios and adjusted p-values, **b)** upset plot for biological pathways, upset plot shows association, and overlapping between different components of a complex system.

**Figure 4:**
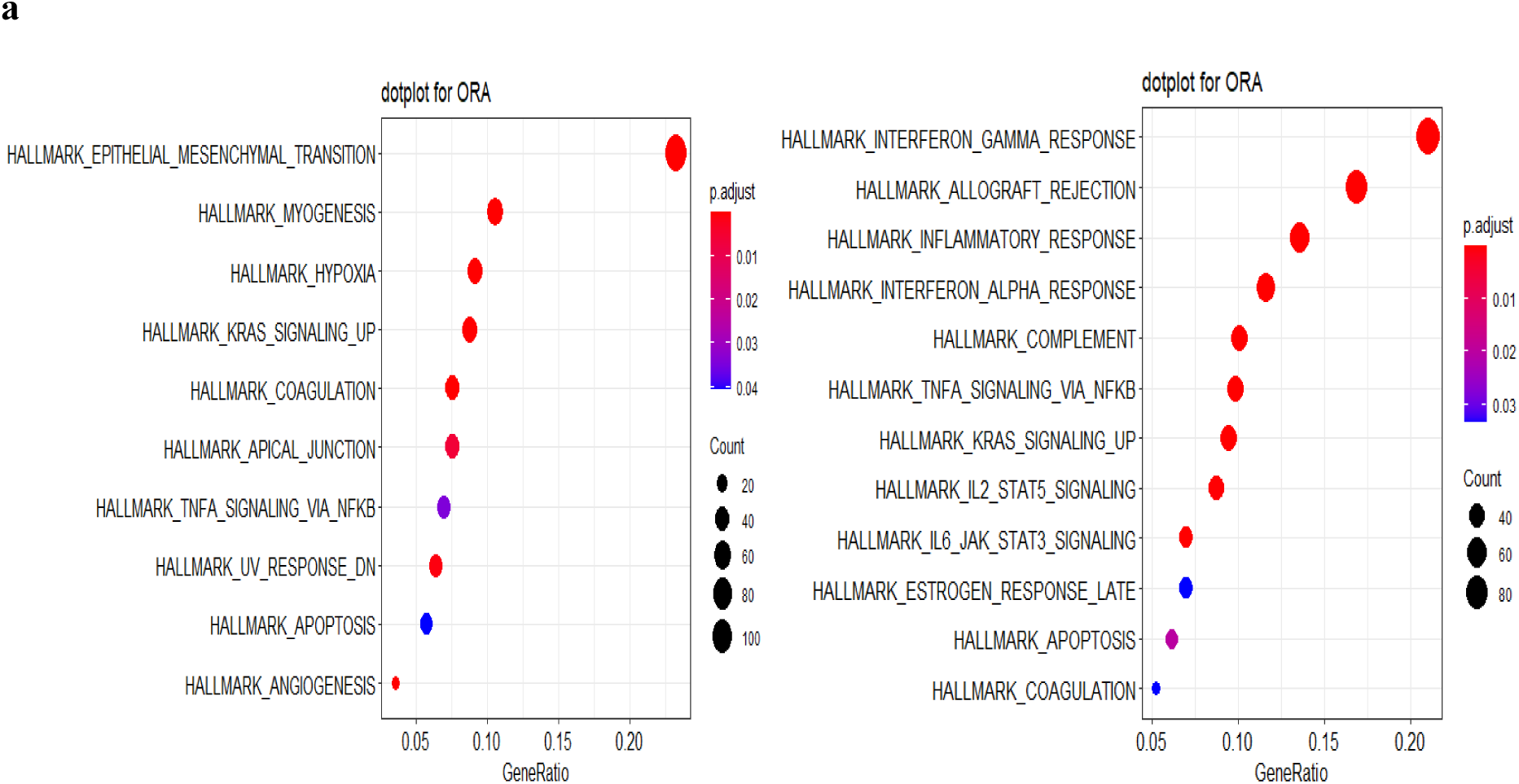
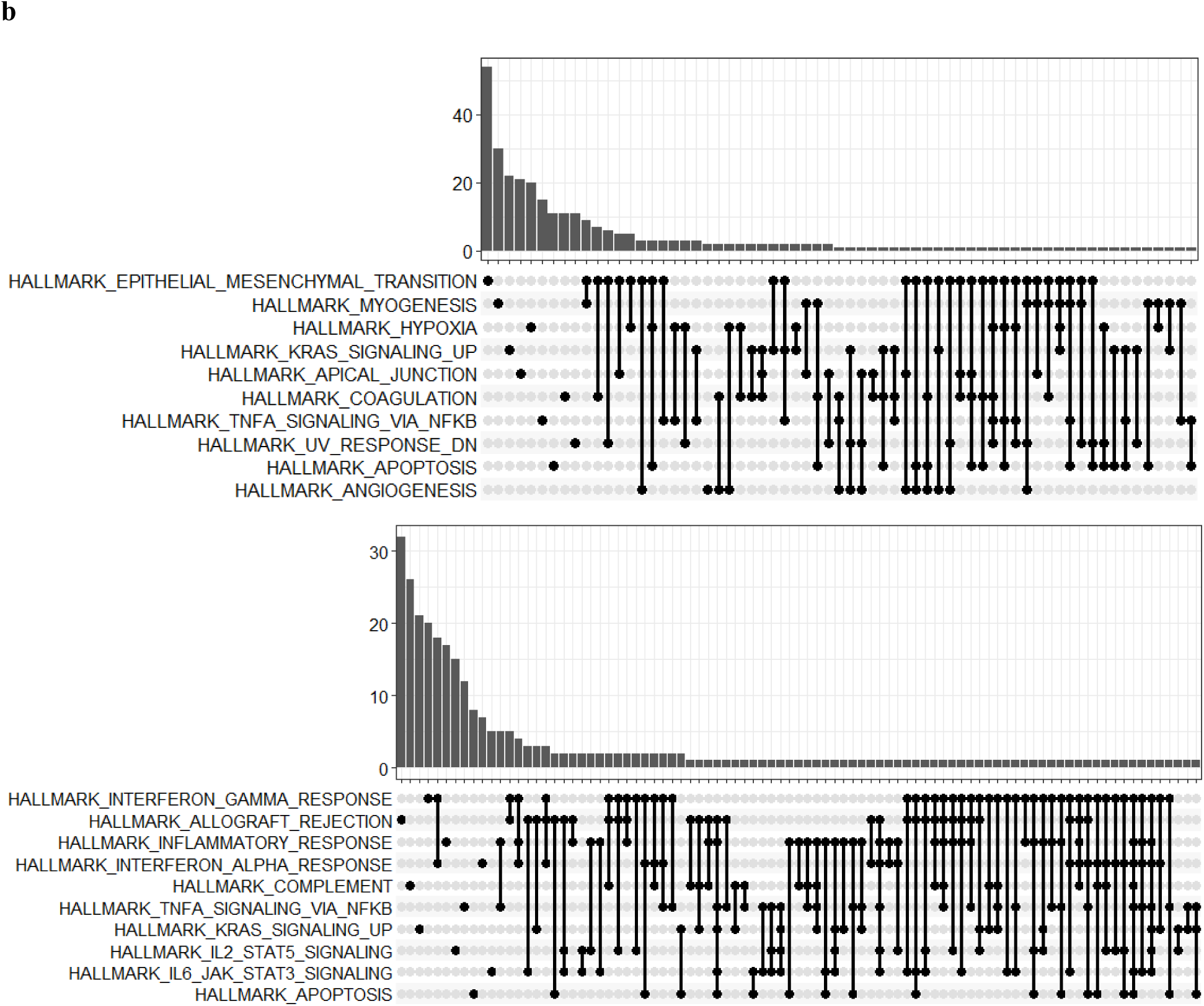
**a)**Dot plot for hallmark analysis, sorted by adjustedP-value, **b)**upset plot for visualizing hallmark analyzing for stage-1 and stage-4 modules

### Identifying key-genes

In order to find key genes in primary modules for either stage, first of all, we constructed a PPI network via STRING. From 1345 genes of four significance modules of stage-1, 941 genes established a network, and from 1276 genes from two statistically meaningful modules of stage-4, 1095 genes were able to assemble a network. We chose eigenvector as a preferred centrality to identify the key genes. Opposed to degree index, which only measures the number of neighbors a node has, eigenvector judges a node based on the significance of its neighbors. According to this concept, if a node connects to more central nodes, it will have a higher value. We chose the top ten genes in the eigenvector category for each stage. Here are some examples of the experimental evidence for some of the key genes. With respect to stage-1 key genes (Figure 5), CD86 gene polymorphism has an association with colorectal cancer risk [35]. ITGAM and ITGAX both could use as potential markers for colorectal cancer [36]. CXCL8 (also known as IL-8) involves in tumorigenesis and progression of CRC. Secretion of CXCL10 and CCL5 by colorectal tumor cells correlate with immune cells infiltration and absence of metastasis [37]. Among the stage-4 key genes (Figure 5), Fibronctine-1 contributes to metastasis in colorectal cancer and has anti-apoptotic effects [38]. COL3A1 also has a role in metastasis and can stimulate proliferation via pi3k-akt axis [39]. MMP9 is a key factor in the reconstruction of ECM, invasion, and regulation of tumor micro-environment, and angiogenesis in colorectal cancer [40]. Furthermore, MMP2 has more than ten times expression in CRCs than healthy tissues. It is one of the prognostic markers of CRC, additionally, it is more involved in local spread than distant metastasis [41].

**Figure 5:**
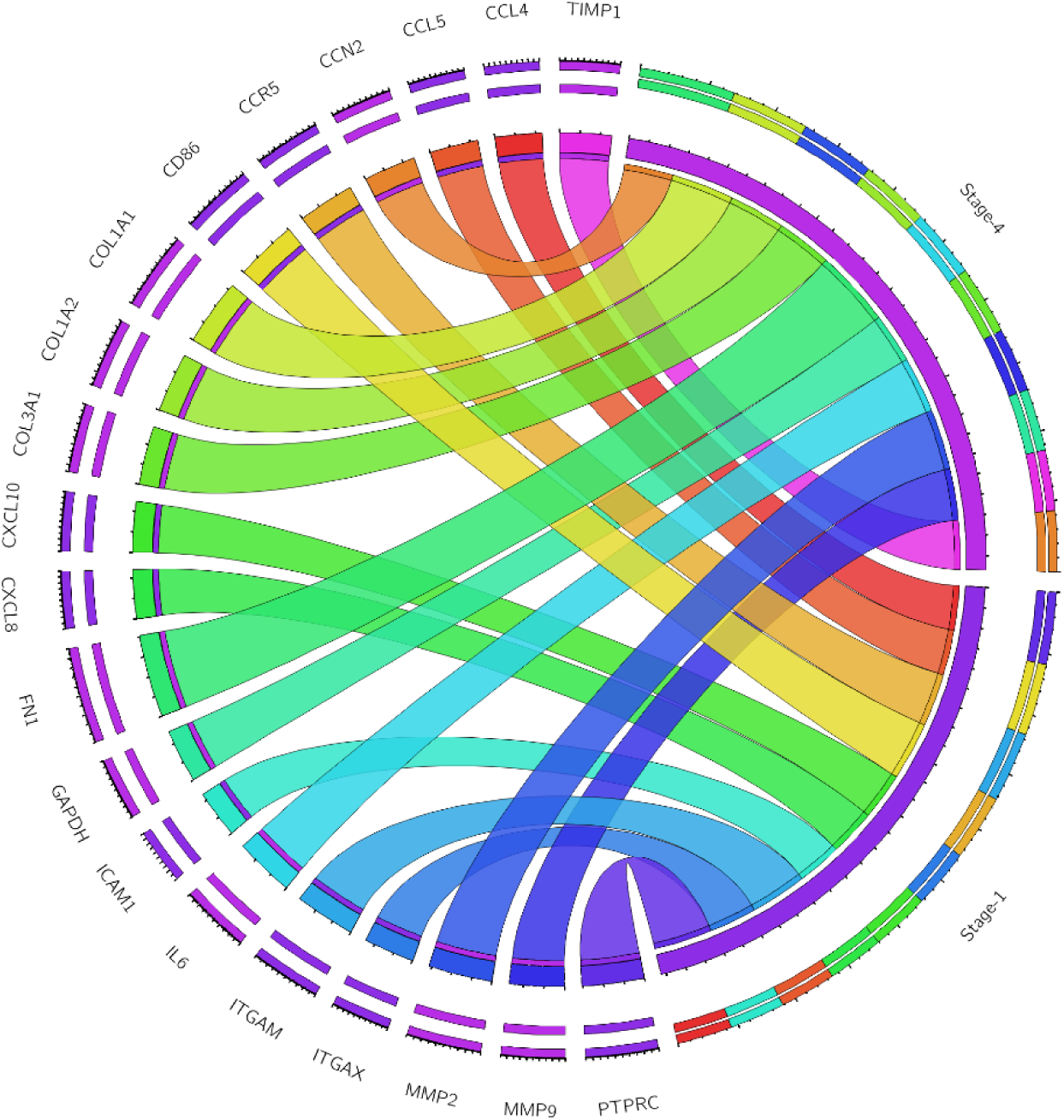
Circus table for genes with the highest eigenvectors’ scores for stage-4 and stage-1. We considered eigenvector as a centrality index in order to identify key genes.

### DEGs results

For DEGs analyze the whole of 25,550 annotated ENTREZ IDs were used. There were 68 samples as stage-1 and 56 samples as Stage-4. The whole study design was stage-4 vs. stage-1 (batch plus condition). Using deseq2 tool, there is no need to remove a batch effect. Instead, it is possible to model a batch effect as a covariate into the analyses during normalization by including batch and condition as a design. Log2fold change was set on 0.75, and adjusted-p value< 0.05. According to the setting, 925 DEGs acquired in which 525 genes were down-regulated (top four of them were L1TD1, CLDN18, DEFA6, and CCL25) (supp2). Also, 400 genes were up-regulated in which HAND1, IGFL1, TMPRSS11E, and FOXG1 were in the top four. This output was used to map the most significant pathway in this process, and finally, we could find out what was the driver path from stage-1 condition to metastatic behavior of late-stage in colorectal cancer (Figure 6).

**Figure 6:**
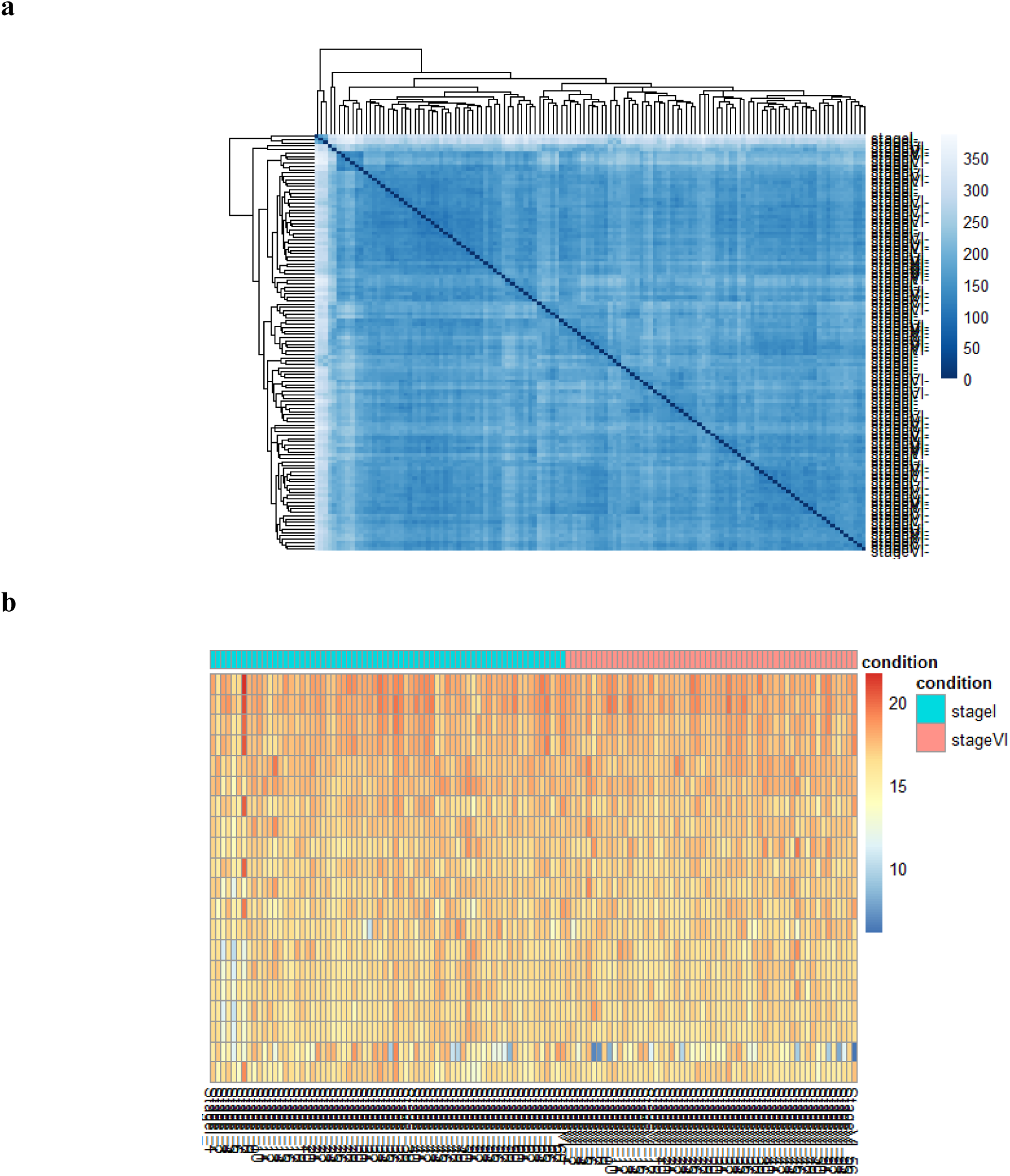
**a)** Heatmaps of sample to sample distance, **b)** Count matrix pheatmap for the DEGs analysis through deseq2 tool.

### Pathway analysis

Based on the results of the differentially expressed gene, we conducted a pre-ranked enrichment analysis via clusterprofiler through wikipathway database. We filtered the DEGs by setting adjusted-p-value <= 0.05 and fold change >= 1.0. Based on the analysis, “LNC in canonical WNT signaling pathway in colorectal cancer” had the best NES (normalized enrichment score) followed by WNT signaling (Figure 7). As it appeared in the analysis, WNT-related pathways have a vital role in the progression from stage-1 to stage-4 of colorectal cancer. Nonetheless, we had two of the most important WNT-LNC related genes: HOTAIR and H19. Since it has already proven by experimental works, WNT signaling might be contributed to tumorigenesis and metastasis in colorectal cancer [42]. Also, it has recently proven that long non-coding RNAs play a role in metastasis of colorectal cancer by modulating WNT signaling pathway [43, 44]. Further, Pi3k-AKT-mTOR pathway also came out as meaningful, especially because of the role that they play in the focal adhesion.

**Figure 7:**
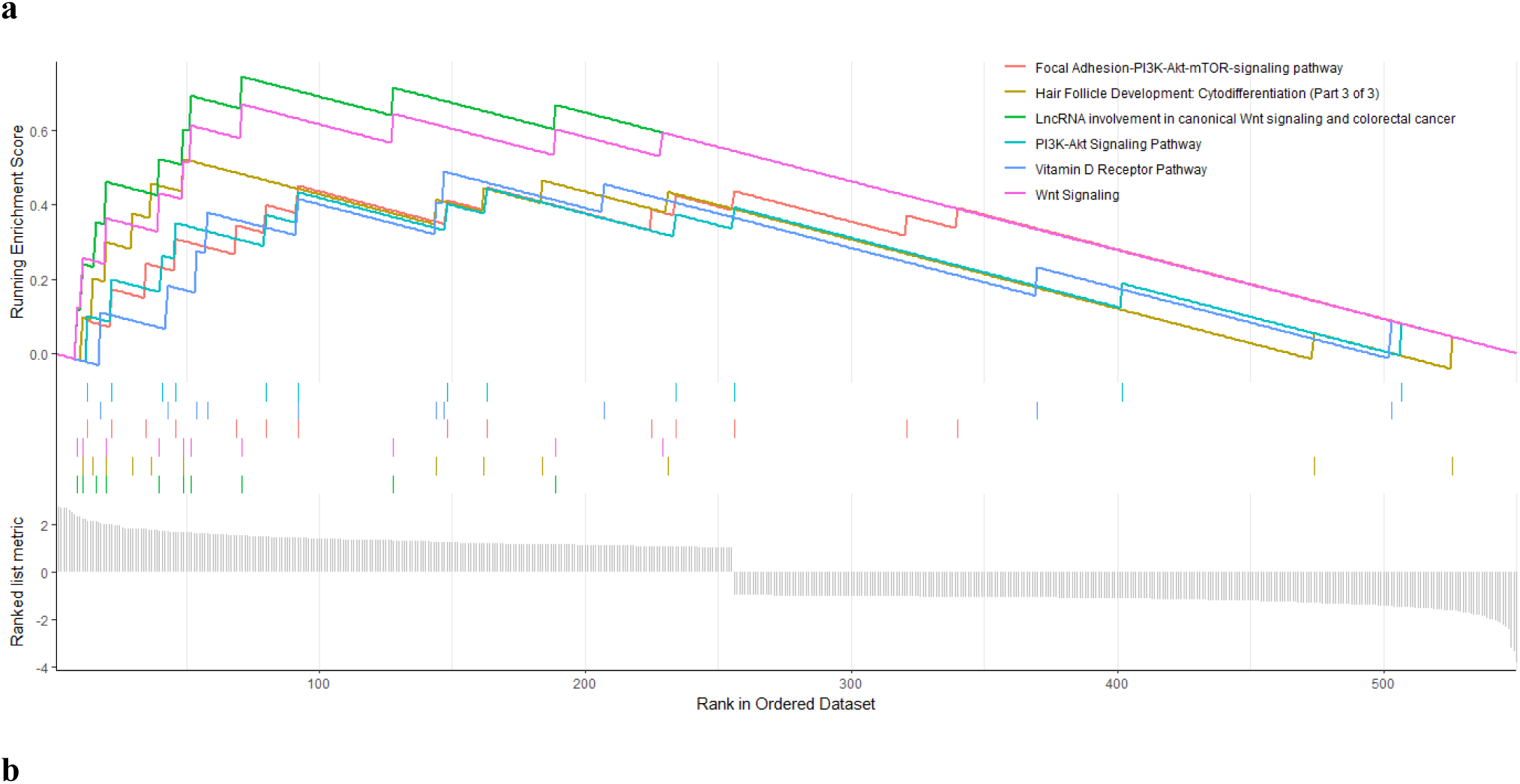
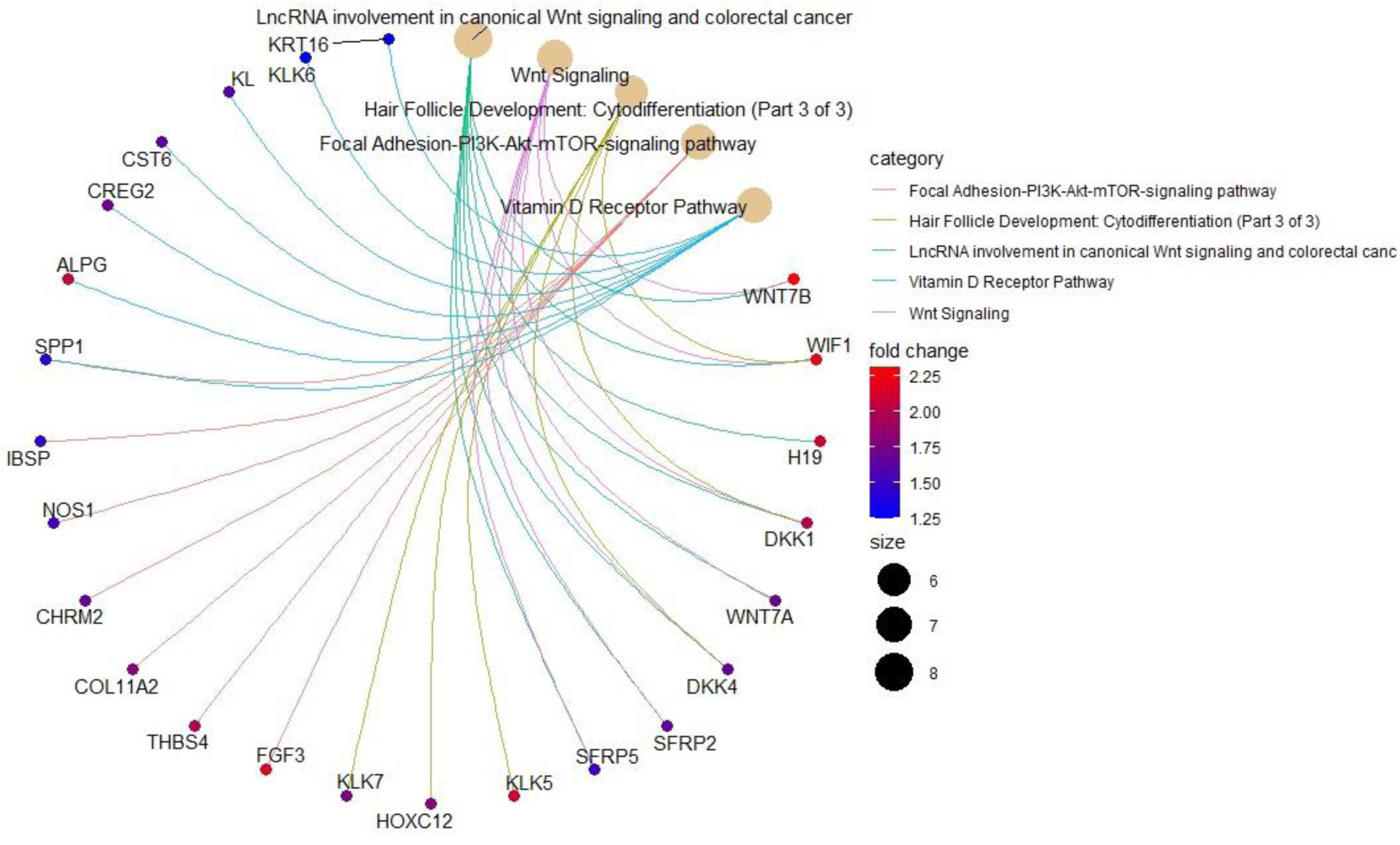
**a)** GSEA plot for our pathway analysis, based on the results, WNT-related pathways were at the outmost of important, **b)** Cnetplot to visualize pathway analysis based on gene-gene interactions and overlapping pathways components

## Discussion

Despite many milestones in treating cancer, still, stage-4 cancers have been problematic. Many cases diagnosed while they have progressed to stage-4, but the disappointment is when a patient shows a complete response to treatment and enters the disease-free survival. Still, after years, cancer will eventually recur, this time more aggressive, more resilient, and even more advanced. It seems it is necessary to have a deep understanding of the underlying mechanism in which cancer progress from stage-1 to stage-4. Knowing the pathways which have direct or indirect involvement in this process could help us to find better and more efficient treatments. In this study, we tried to identify significance modules in the initial and late stages of colorectal cancer, and then analyze them accordingly. Based on the results, an immune-related process, specifically interferon-gamma response, has a distinct role in stage-1 colon adenocarcinoma, but when it comes to stage-4, the pathways and processes are linked to extracellular matrix and invasion. We also performed a topological analysis on the networks obtained from these modules and identified PTPRC, CD86, ITGAM, CCR5, and CXCL10 as the top five critical genes in stage-1, whereas FN1, COL1A1, MMP2, COL1A2, and IL6 were identified as the top five in stage-4 of adenocarcinoma. Interestingly, interferon gamma (IFN-y) uses as a treatment for the colorectal cancer and also many other forms of malignancies, since it is one the essential component of the TH-1 response [45]. But recent researches found out that there could be more to tell about the role of IFN-y in tumor and tumor microenvironment. In one study, researchers concluded that low-dose of IFN-y could contribute to cancer stemness and metastasis and its effects are dose-dependent. Also there is some reports about contribution of IFN-y in aggressiveness and proliferation of tumor cells [46, 47].

According to the differentially expressed gene analysis, the WNT signaling and LNC in canonical WNT signaling in colorectal cancer might be one of the main reasons for the malign transition from stage-1 to stage-4. It is clear that WNT signaling is one the most frequently hyperactivated path in all types of colon cancers, But the most types of CRCs have WNT ligands-independent hyperactivity, which means direct neutralization of WNT-ligands cannot be effective [48]. Nonetheless, in our analysis, not only the signaling itself but also the WNT-ligands were up-regulated. Still worth of mention that according to the previous studies, WNT signaling might be responsible for checkpoint inhibitor resistance and immunological difficulties in treating colorectal cancer [48–50]. As it appear in our study, lncRNA in canonical WNT signaling in colorectal cancer also has a significant role in the transition from stage-1 to stage-4; this notion additionally has been proven by experimental works [43, 51]. HOTAIR is one of these LNCs that could contribute to the overexpression of WNT signaling [52]. HOTAIR has a role in metastasis, drug resistance, and proliferation [53]. According to our DEGs analysis, HOTAIR is one of the most expressed differentially from the stage-1 to stage-4. H19 is another related LNC in WNT signaling in metastasis of colorectal cancer. H19 has a role in metastasis, proliferation, and drug resistance in many types of cancers [54]. Increased expression of H19, also correlates with poor prognosis [55]. H19 has many important pathways under its control like MYC and let-7, methylation of the genome, regulation of cell cycle, and of course WNT-Beta catenin [56–58]. In our DEGs analysis, the expression of H19, was meaningfully increased.

## Conclusion

In the past studies, it has been shown that inhibition of HOTAIR or H19 could significantly reduce invasion, proliferation, and drug resistance in colorectal cancer [59, 60]. Now, our systems biology study could systematically support the idea that the WNT-signaling and LNCs mediated in WNT signaling, specifically HOTAIR and H19, might have a notable effect in the metastatic process in colorectal cancer. Therefore their inhibition could be considered as a therapeutic option to prevent the metastasis and progression of colorectal cancer.

## Declarations

### Ethics approval

This study does not need any ethical approvals or informed consent.

### Consent for publication

Not Applicable.

### Supplementary data and materials

Supplementary file 1: Phenotypes and samples information

Supplementary file 2: Deg results

### Code availability

All software have been mentioned in the manuscript and all scripts would be available upon request.

### Authors’ Contributions

MG designed the study, generated and analyzed the data, and drafted the manuscript. YA supervised the study, validated the data, and reviewed the manuscript. All authors read and approved the final manuscript.

## Acknowledgments

The authors acknowledge the reviewers for their helpful comments.

